# The *Drosophila melanogaster* Y-linked gene, *WDY*, is required for sperm to swim in the female reproductive tract

**DOI:** 10.1101/2023.02.02.526876

**Authors:** Yassi Hafezi, Arsen Omurzakov, Jolie A. Carlisle, Ian V. Caldas, Mariana F. Wolfner, Andrew G. Clark

## Abstract

Unique patterns of inheritance and selection on Y chromosomes lead to the evolution of specialized gene functions. Yet characterizing the function of genes on Y chromosomes is notoriously difficult. We report CRISPR mutants in Drosophila of the Y-linked gene, *WDY*, which is required for male fertility. *WDY* mutants produce mature sperm with beating tails that can be transferred to females but fail to enter the female sperm storage organs. We demonstrate that the sperm tails of *WDY* mutants beat approximately half as fast as wild-type sperm’s and that the mutant sperm do not propel themselves within the male ejaculatory duct or female reproductive tract (RT). These specific motility defects likely cause the sperm storage defect and sterility of the mutants. Regional and genotype-dependent differences in sperm motility suggest that sperm tail beating and propulsion do not always correlate. Furthermore, we find significant differences in the hydrophobicity of key residues of a putative calcium-binding domain between orthologs of *WDY* that are Y-linked and those that are autosomal. Given that *WDY* appears to be evolving under positive selection, our results suggest that *WDY*’s functional evolution coincides with its transition from autosomal to Y-linked in *Drosophila melanogaster* and its most closely related species. Finally, we show that mutants for another Y-linked gene, *PRY*, also show a sperm storage defect that may explain their subfertility. In contrast to *WDY, PRY* mutants do swim in the female RT, suggesting they are defective in yet another mode of motility, navigation, or a necessary interaction with the female RT. Overall, we provide direct evidence for the long-held presumption that protein-coding genes on the Drosophila Y regulate sperm motility.

## INTRODUCTION

Y chromosomes are unique in the genome of many organisms, including mammals and Drosophila, in being haploid, male-limited, repeat-rich, highly heterochromatic, and, in particular, having reduced or no recombination ^1^. The resulting selective pressures on Y chromosomes cause rapid degeneration of most protein-coding genes, yet a few genes are maintained on Y chromosomes with remarkable evolutionary endurance. Such genes are maintained for extended periods under strong purifying or sometimes positive selection, repeatedly and independently acquired in different lineages, or undergo massive copy-number amplification on the Y chromosome ^2–5^. These patterns of variation indicate that selection favors placing such genes into this seemingly inhospitable genomic environment. In support of this concept, there is striking similarity in both the expression patterns and functions of many Y-linked genes ^6–8^.

The 40 MB *Drosophila melanogaster* Y chromosome contains only 14 known protein-coding genes ^9–11^. X0 flies are male yet sterile – therefore, the Y chromosome is required for male fertility but not sex determination or viability ^12^. Six genetic loci on the Y, known as the “fertility factors”, contribute to this fertility function. The fertility factors were defined by a series of X-ray induced X-Y translocations ^13,14^ and, remarkably, half of them were discovered to be axonemal dyneins ^15,16^, suggesting that the *Drosophila melanogaster* Y chromosome plays a pivotal role in sperm motility. *kl-1* males, in contrast to all other fertility factor mutations, produced mature and motile sperm despite being completely sterile^17^. *kl-1* sperm were transferred to the female reproductive tract (RT) following mating but could not be recovered from the female sperm storage organs. The specific defect that prevents *kl-1* mutant sperm from entering storage or fertilizing eggs is unknown. The molecular identity of *kl-1* remained unknown until, recently, the gene, *WDY*, was found to be contained within the *kl-1* region ^18^ and required for male fertility based on RNA interference (RNAi) ^19^. Other protein-coding genes or functional repetitive elements may still reside in the *kl-1* genetic region, which was estimated cytologically to span ∼3% of the length of the Y chromosome ^20^, and it is unclear whether *WDY* mutants produce mature sperm or show a sperm storage defect. More generally, the importance of motility for sperm storage and the mechanisms that regulate sperm motility remain poorly understood in Drosophila.

Here we generated CRISPR mutants to investigate the function of *WDY*. We demonstrate that *WDY* mutant sperm display the storage defect suggested for *kl-1*. Furthermore, mutant sperm have reduced beat frequency and are unable to swim beyond the seminal vesicle. We show that mutants in another Y-linked gene, *PRY*, also have impaired sperm-storage. *WDY* and *PRY* are both evolving under positive selection^2^ and Y-linked orthologs of *WDY* show significant changes in key amino acid residues in a conserved calcium-binding domain, suggesting directed functional evolution of this gene.

A high incidence of genes with predicted sperm motility functions are seen on Y chromosomes across many species from Drosophila to great apes ^7,21^. Carvalho et al (2000) hypothesized that, in species where there is a high level of sperm competition (such as *Drosophila melanogaster*), motor proteins are specifically recruited to the Y chromosome where they can evolve without constraint from male-female antagonistic selective forces. Our study provides an in-road to studying the evolutionary logic of this association.

## RESULTS AND DISCUSSION

### *WDY* mutants are sterile but produce mature, motile sperm

We used CRISPR to precisely target *WDY*. One of the major challenges of studying the Y chromosome is in propagating sterile mutations on a haploid chromosome. We used a crossing strategy involving compound sex chromosomes to make and stably propagate heritable mutations in *WDY* (mutant stocks consist of 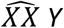,*WDY* females and 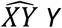,*WDY* males, see Methods, Figure S3 ^19^). Our crossing scheme also enabled us to identify and eliminate large chromosomal truncations that are common during genetic editing of the Y chromosome, likely due to its highly repetitive nature ^19^. To evaluate the phenotype of our mutants, we then removed the compound chromosomes by selective breeding to generate X Y, *WDY* mutant males. We confirmed each phenotype with three different *WDY* alleles, *F8, C104*, and *C3*, containing deletions of 547 bp, 545 bp and 443 bp, respectively (Figure S4, Table S4). All are large deletions close to the N-terminus that disrupt the reading frame and are therefore expected to be null; all three gave the same phenotype. We compared mutants to controls that account for the genetic background (Y^Tomato^) or crossing scheme (Y^C7^ and Y^G107^). In individual crosses to females from a wild-type strain (Canton S), control males produced progeny while *WDY* males were sterile (Table S5).

To investigate the cause of this sterility, we first examined the distribution of sperm in the testes using Protamine-GFP ^22^, which labels sperm heads. Sperm of mutants in Y-linked fertility factors kl-2, kl-3,kl-5, ks-1, and ks-2 are eliminated before this time, during the individualization stage^19,23,24^. In contrast, we observed an accumulation of *WDY* mutant sperm in the posterior-most section of the testes, where individualized sperm accumulate while sperm coiling occurs, causing that region to bulge in the mutant (Figure 1). Sperm coiling is thought to function as a quality control step during which sperm with abnormal tails are eliminated by ingestion by the terminal epithelium ^25^. The accumulation of *WDY* sperm in the posterior testes may be due to their progression being stalled by this quality control mechanism or may indicate insufficient motility to exit the testes.

**Figure 1.**
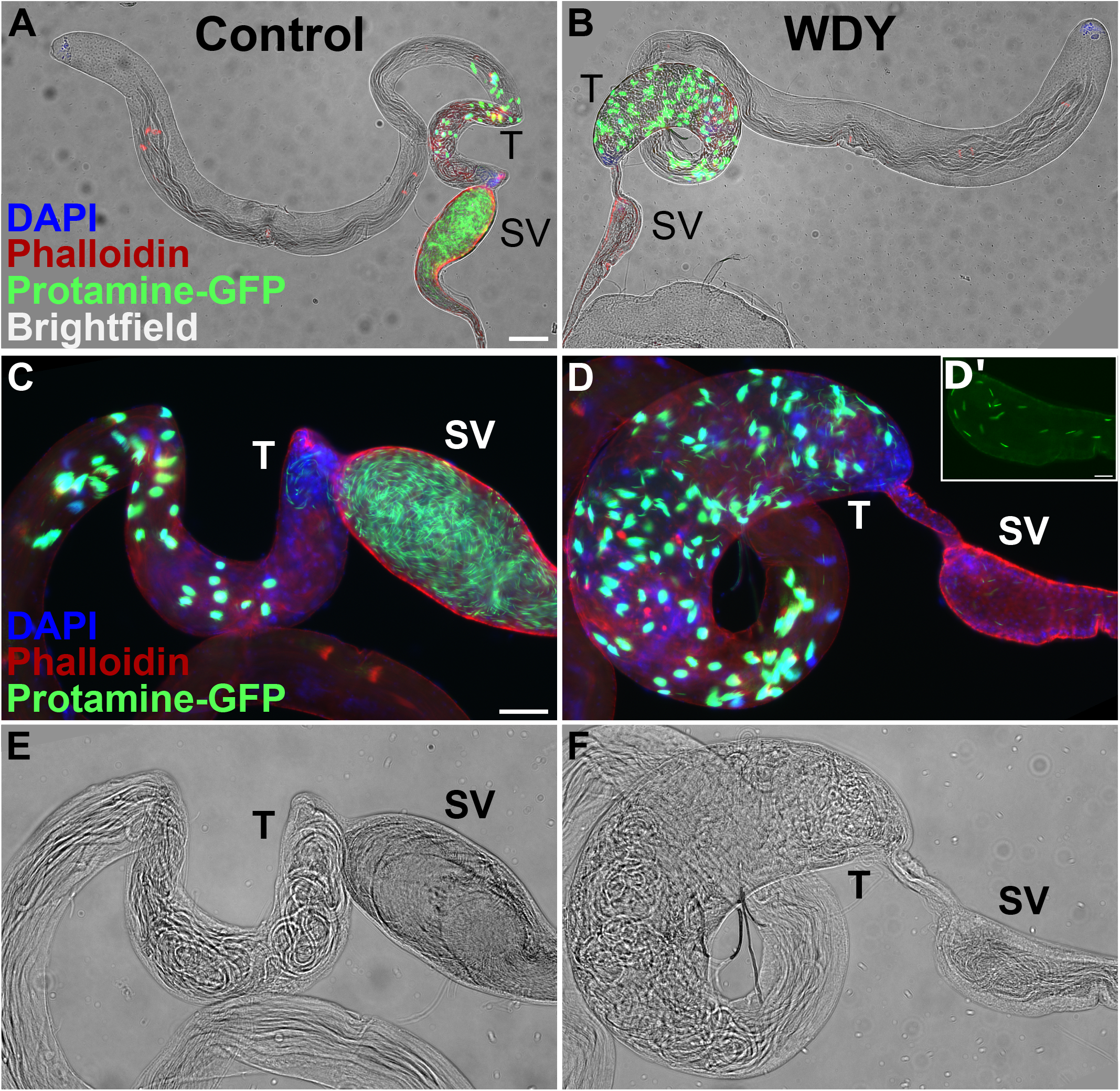
Spermatogenesis is backed up in *WDY* testes, but some mature sperm are found in the seminal vesicle. Whole testes from control (A) and *WDY* mutant (B) males whose sperm were labelled with Protamine-GFP (green), Phalloidin (red) and DAPI (blue) overlaid on brightfield images. C-F Higher magnification view of posterior testes and seminal vesicle. Inset (D’) shows Protamine-labelled sperm in seminal vesicle of mutant. Testes (T) and seminal vesicle (SV) are marked. Bar denotes 100 µm for A-B, 50 µm for C-F, and 20 µm for D’.

In the seminal vesicles there were fewer sperm in *WDY* mutants than controls (Figure 1C, D). Yet tails of sperm from both *WDY* and control males were observed to beat after we tore open the seminal vesicles (Movie M1,2). We observed no obvious differences in the movement of *WDY* versus control sperm. These observations match Kiefer’s conclusion that *kl-1* mutants were sterile but produced seemingly motile sperm ^17^. Our results that *WDY* mutations are sufficient to result in sterility, yet produce sperm that are motile, make it highly likely that *WDY* is the fertility factor known in the literature as *kl-1*.

### *WDY* mutant sperm are transferred to females, but do not enter the storage organs

We next tracked the movement of Protamine-labelled sperm in the RT of wild-type (Canton S) females 30 min after the start of mating (mASM). Sperm from both control and *WDY* mutant males were found in the female’s uterus (bursa) (Figure S5) and their tails were observed to beat when dissected out of the uterus (Movie M3,4). In both *WDY* and control genotypes, an open or folded conformation of the uterus correlated with the presence or absence of sperm, respectively, as expected ^26,27^. We conclude that motile *WDY* sperm are transferred to females and that *WDY* seminal fluid induces conformational changes in the uterus.

We did, however, observe defects in the number of sperm transferred to, and the distribution of the sperm within, the female RT. *WDY* males transferred less than half as many sperm as control males as quantified at 30 mASM (Figure 2A, p<0.001). After mating, Drosophila sperm move rapidly from the uterus into either the primary storage organ, the seminal receptacle, or one of two long-term storage organs, the spermathecae ^28^ (Figure 2B). At 30 mASM most control samples contained some stored sperm, but no *WDY* sperm were found in the storage organs. At 2 hASM maximal numbers of sperm are stored in most control samples ^29^. Yet, again, no sperm from *WDY* mutants were seen in storage (Figure 2C-E). We also examined sperm in RTs of females left to mate overnight to see if a longer time or multiple mating may enable sperm to enter storage (Figure S5). Control samples all had stored sperm. *WDY* sperm were regularly observed in the uterus but never in any of the storage organs. We conclude that *WDY* mutant sperm are unable to enter the storage organs. In many animals, storage is required for sperm to become competent for fertilization ^30,31^, thus the lack of sperm storage might explain why *WDY* males are sterile.

**Figure 2.**
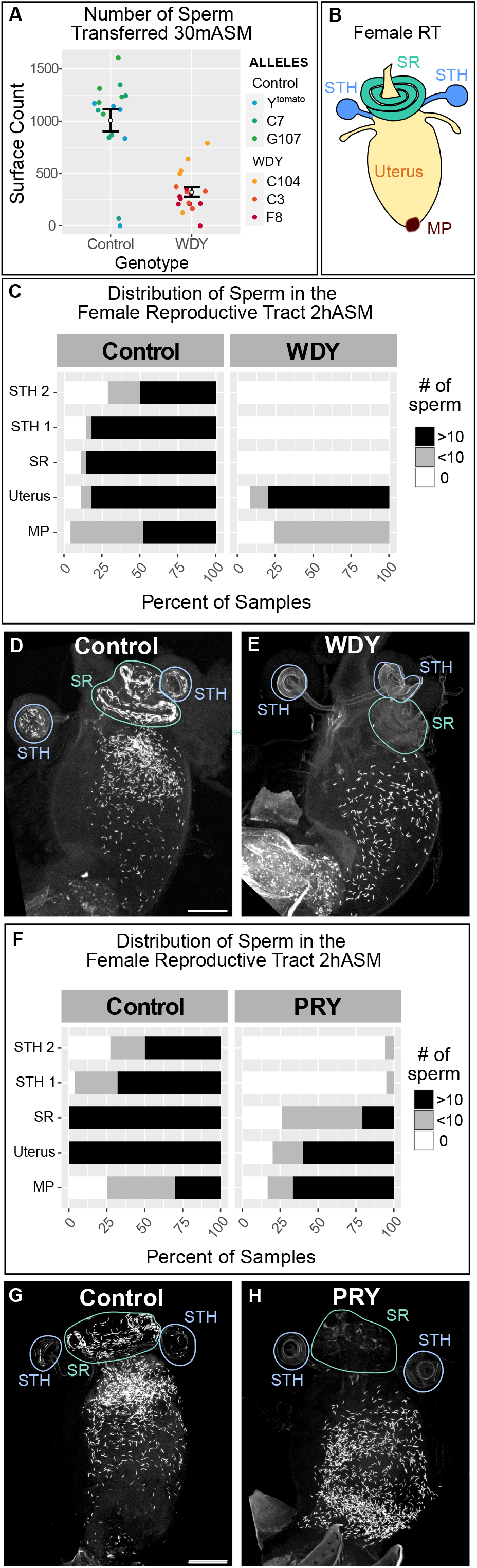
*WDY* and *PRY* mutant sperm fail to enter the storage organs in the female RT. (A) Quantification of the number of sperm transferred to the female uterus 30 mASM to control or *WDY* mutant males. (B) Cartoon of the female RT indicating the mating plug (MP, brown), uterus (yellow), and the storage organs – the seminal receptacle (SR, green) and two spermathecae (STH, blue). (C) Quantification of the distribution of sperm within the female RT 2hASM. Representative images show Protamine-labelled sperm from Control (D) and *WDY* (E) mutant males within the female RT 2 hASM. (F) Quantification of the distribution of sperm within the female RT 2 hASM. Representative images show Protamine-labelled sperm from Control (G) and *PRY* (H) mutant males within the female RT 2 hASM. Bar denotes 100 µm.

### *WDY* mutant sperm in the male seminal vesicle and female uterus have decreased beat frequency

Although *WDY* mutant sperm beat visibly in vitro (Movie M1-4), we wished to test whether subtle motility defects prevent them from being able to enter storage. We measured the beat frequency of sperm tails, by recording videos of control and *WDY* mutant sperm dissected directly from the male seminal vesicle or from the female uterus at 30 mASM (Methods, Movie M1-4, Figure 3A, B). In the seminal vesicle, the fastest *WDY* mutant sperm tails beat at an average of 6.0 Hz, whereas the fastest tail-beats by control sperm average 12.3 Hz (p<0.001). In the female uterus, the fastest *WDY* mutant sperm tails beat an average of 7.0 Hz, whereas the fastest control sperm tails beat at an average of 13.1 Hz (p<0.001). We conclude that *WDY* mutant sperm have a lower tail-beat frequency than wild type sperm in both the male and female RTs.

**Figure 3.**
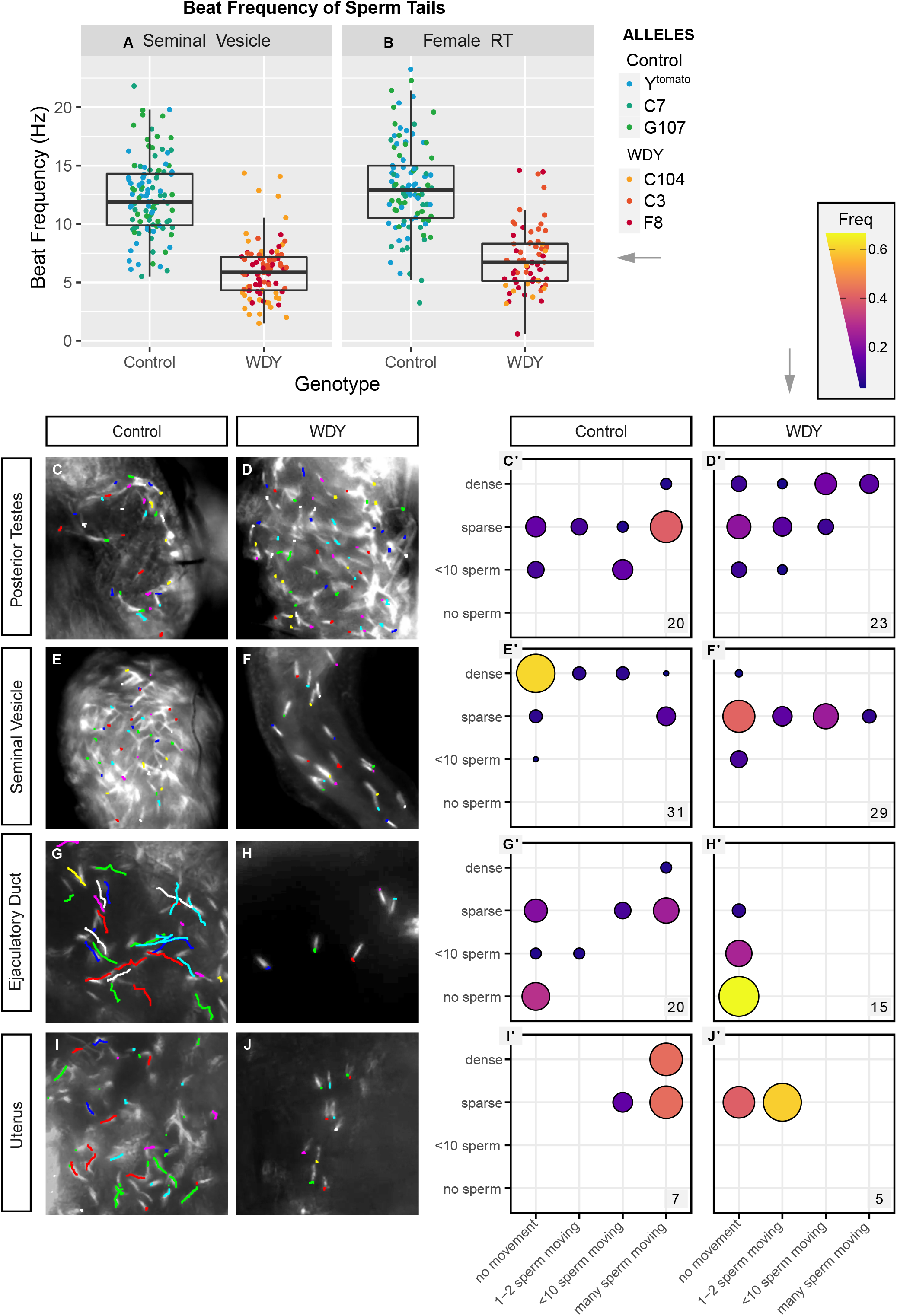
*WDY* mutant sperm have reduced beating frequency and do not swim in the female RT. Quantification of tail beating frequency of *WDY* and control sperm dissected from seminal vesicles (A) and the female RT 30 mASM (B). Manual tracking of Protamine-GFP-labelled control (C,E,G,I) and *WDY* mutant (D,F,H,J) sperm heads over a 0.5 sec interval for representative videos of the posterior testes (C,D), seminal vesicle (E,F), ejaculatory duct (G,H), and uterus 1h ASM (I,J). (C’-J’) Corresponding quantification of the number of sperm heads and degree of movement observed from videos of each region of the RTs. The number of each type of organ that was scored is indicated.

### *WDY* mutant sperm are unable to swim in the male ejaculatory duct and female uterus

We hypothesized that the lower beat frequency affects the ability of sperm to propel themselves. To test for defects in sperm swimming, sperm movement was assessed in videos by tracking the Protamine-labelled heads of control and *WDY* mutant sperm. In all regions the swimming speed of individual sperm varied, but there was an overriding regional pattern to the motility (Figure 3C-J). In mammals, sperm leaving the testes are immotile and must go undergo “epididymal maturation” in order to gain the ability to move progressively and to fertilize eggs ^32^. It was previously suggested, and often repeated, that, similarly, Drosophila sperm do not gain motility until they reach the seminal vesicle ^33^. We were surprised to see some individualized sperm heads slowly swimming within the posterior testes of most control samples (Figure 3C,C’, Movie M5). This suggests that in Drosophila sperm motility is normally initiated within the testes. *WDY* mutant sperm heads in this region also often moved around, suggesting that at least some mutant sperm develop motility (Figure 3D,D’, Movie M6).

Individual sperm heads generally ceased to move in the seminal vesicles of both control and *WDY* mutant flies, while flagella remained beating. However, mass movements occurred from contractions of the whole organ. It was unclear whether dense packing of sperm or some physical or chemical property of the seminal vesicle caused the immobilization of sperm heads while sperm tails continued to beat vigorously (Figure 3E,E’, Movie M7). This highlights that flagellar beating does not necessarily correlate with sperm swimming (i.e. moving through space). There were far fewer sperm in *WDY* mutant seminal vesicles, but the mutant sperm heads were predominantly immobilized, as in controls (Figure 3F,F’, Movie M8). We conclude that at least some *WDY* sperm develop the ability to swim in the testes and become immobilized in the seminal vesicle, as normal.

In contrast, a striking difference was seen in the ability of *WDY* sperm to swim beyond the seminal vesicle. In samples where sperm were found in the ejaculatory duct, control sperm heads were observed to move swiftly while *WDY* sperm heads appeared motionless (Figure 3G-H, G’-H’, Movie M9-10), and the same pattern was observed for sperm heads in the uterus 1 hASM. These sperm appear to be alive, as the tails continue to beat in place (Figure 3A,B). The lack of swimming *WDY* sperm in the uterus likely explains the inability of *WDY* sperm to enter the storage organs, though other defects may also exist and contribute ^34^. That *WDY* sperm in the posterior testes can swim suggests either (1) there is a subclass of *WDY* sperm that are capable of swimming but degenerate or (2) *WDY* mutant sperm are unable to navigate between different regions of the RTs.

### Significant hydrophobicity differences in putative calcium-binding residues coincide with *WDY*’s transition to Y-linkage in *Drosophila melanogaster*

Calcium regulates sperm motility in many organisms, including humans ^39^ and Drosophila ^40–42^. WDY’s amino acid sequence contains a calcium-binding domain signature: an EF Hand (Interpro ^36^). Functional EF Hand domains contain a pair of motifs, each consisting of a loop flanked by alpha helices, that can bind Ca^2+^ ions. The specific characteristics of the loop affect calcium-binding affinity ^37^. We identified a putative pseudo EF hand motif followed by a canonical EF hand motif in WDY (Methods, Figure 4A-C). Known calcium-binding proteins (e.g. Calbindin D9K ^38^) also display this configuration. We also improved the annotation of two WD40 domains (Methods, Figure 4A), which typically mediate protein-protein interactions in protein complex assembly and/or signal transduction. Based on these findings, we speculate that *WDY* is necessary for sperm to recognize and adjust their motility based on differences in calcium in different regions of the RT.

**Figure 4.**
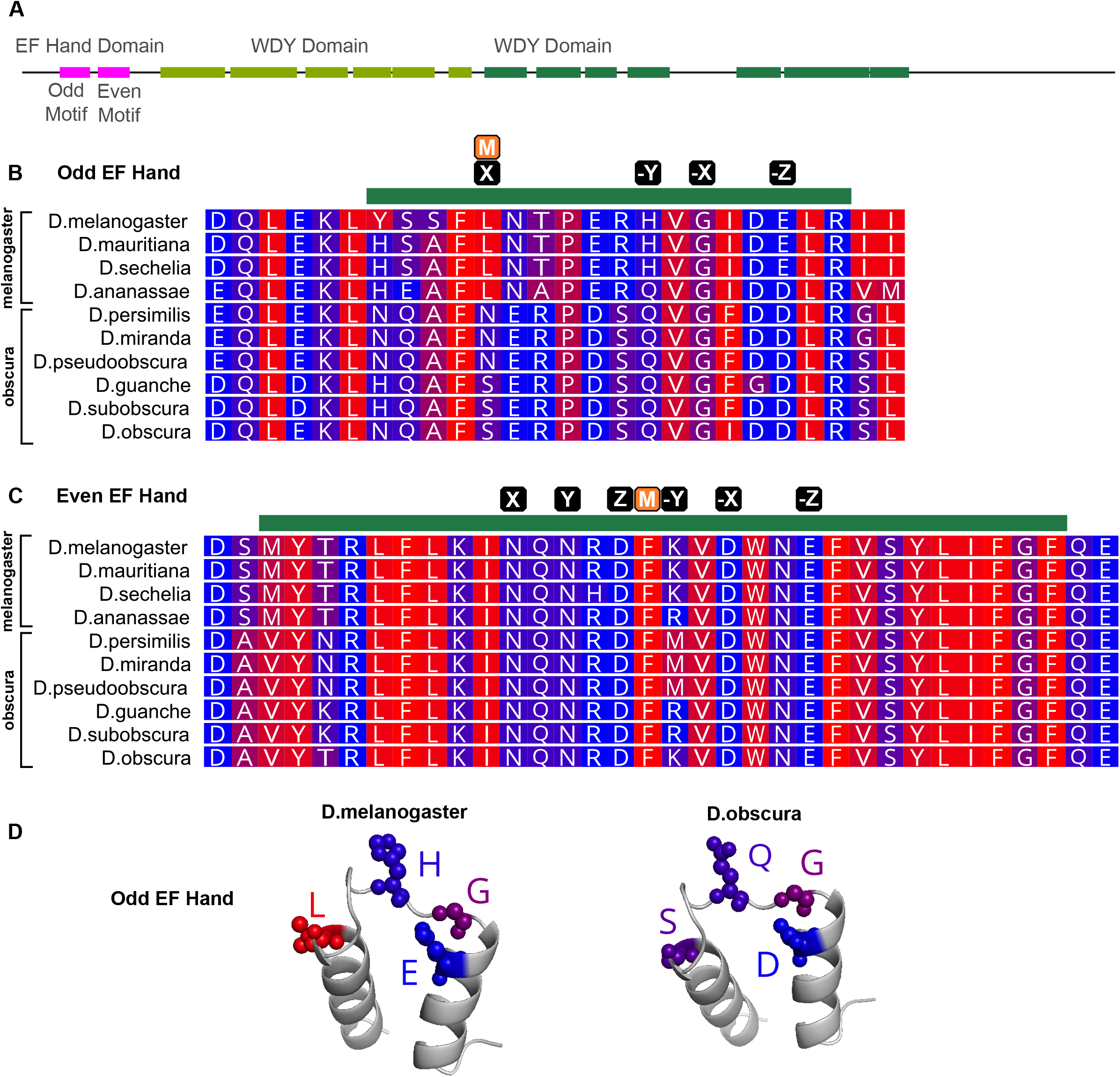
Difference in hydrophobicity in the EF Hand Domain in Y-linked orthologues of *WDY* in *Drosophila melanogaster* and its closest relatives. (A) Domain structure of WDY (B,C) Protein alignment of melanogaster and obscura group species for the region with the Odd (Pseudo) (B) and Even (Canonical) (C) EF Hand motif. Green bar indicates the motif, M indicates the position of any mismatch between the *melanogaster* sequence and the consensus, and black boxes indicate putative calcium-binding residues (X,Y,Z,-X,-Y,-Z). (D) Predicted AlphaFold structure of Odd EF Hand domain with putative calcium binding residues labelled, generated in PyMOL. Blue-red scale indicates hydrophobicity (Red is hydrophobic, Blue is hydrophilic).

We also compared sequences of the EF Hand domain between WDY orthologs in the melanogaster species group, where WDY is Y-linked, and the obscura species group, where WDY homologs are autosomal or X-linked ^4^. While there was relative conservation of the domain within each group (82.4% identical sites in each group), there were notable differences between the groups (63.2% identical sites overall), particularly in the loop residues of the Odd EF Hand motif (Figure 4B). The transition at position “X” is particularly compelling, since it involves a profound biochemical change in a conserved residue thought to directly bind calcium ^38^. The shift away from canonical residues in the melanogaster group could indicate a modulation of calcium binding, and thus significant functional evolution in the EF Hand domain, coinciding with Y-linkage.

### *PRY* is also required for efficient sperm storage

We previously generated and characterized mutants in another Y-linked gene, *PRY*, whose phenotype was consistent with abnormal sperm storage in females: mutants had low levels of fertility on the first day after mating but no fertility on subsequent days ^19^. Our finding that *WDY* affects sperm entry to the storage organs led us to wonder whether *PRY* affects a similar step. In contrast to *WDY, PRY* mutant sperm do swim in the female RT (Movie M13). However, the number of *PRY* sperm stored was significantly reduced compared to controls at 2 hASM (Figure 2F-H). *PRY* mutant sperm were frequently absent or reduced in the seminal receptacle, and rarely observed in the spermathecae. The number of stored *PRY* sperm was similar in RTs 2 hASM and 24 hASM (Figure S5) – defined times after a single, observed mating. However, significantly more sperm entered storage organs if males and females were housed together overnight (Figure S5). Seminal fluid proteins responsible for long term physiological effects of mating on females, including the inhibition of remating, can bind to sperm tails ^2,43–45^. Thus, lack of stored *PRY* sperm may lead to increased remating in these females.

Overall, we present functional evidence for a role for two Y-linked genes in sperm storage and demonstrate that *WDY* mutants have specific defects in sperm motility. Two broad themes emerge from this work. First, sperm may be unable to enter storage for different reasons – insufficient swimming speeds or an inability to navigate or gain entry into the storage organs. We demonstrate that sperm motility defects do manifest in an inability to enter storage. Female secretions are necessary to promote sperm storage^46^ so entry to sperm storage may be a hurdle imposed by females to ensure that only sperm with a certain level of motility/fitness are able to fertilize eggs. Higher remating rates upon lower sperm storage, as we appear to see with *PRY*, would then likely contribute to elimination of such sperm from the female RT through sperm competition.

Second, across species Y-linked genes appear to show ‘functional coherence’ ^6,23^. Even within the realm of male fertility, a disproportionate number of Y-linked genes seem to be singularly focused on aspects of sperm motility ^15,16^. Three axonemal dyneins were previously discovered on the Drosophila Y chromosome ^15,16^. However, since no sperm are produced upon genetic ablation of five out of the six fertility factor genes, the role of these genes in sperm motility has never been observed ^13,14,19,23^. We now add strength and stringency to the picture of functional coherence on the Drosophila Y chromosome. On the one hand being Y-linked allows sperm motility genes to escape the problems of countervailing selection in females (sexual conflict). However, being Y-linked bears the cost of not being able to recombine, which reduces the efficacy of natural selection (the Hill-Robertson effect). The fact that so many sperm motility genes are retained on the Y chromosome indicates a dynamic balance between these two opposing selective forces in different regions of the genome.

## FIGURE LEGENDS

**Movie M1: Sperm dissected from seminal vesicles of control males**.

**Movie M2: Sperm dissected from seminal vesicles of *WDY* males**.

**Movie M3: Sperm dissected from the uterus of females mated to control males**.

**Movie M4: Sperm dissected from the uterus of females mated to *WDY* males**.

**Movie M5: Control sperm heads in the posterior testes**.

**Movie M6: *WDY* sperm heads in the posterior testes**.

**Movie M7: Control sperm heads in the seminal vesicle**.

**Movie M8: *WDY* sperm heads in the seminal vesicle**.

**Movie M9: Control sperm heads in the ejaculatory duct**.

**Movie M10: *WDY* sperm heads in the ejaculatory duct**.

**Movie M11: Control sperm heads in the uterus**.

**Movie M12: *WDY* sperm heads in the uterus**.

**Movie M13: *PRY* sperm heads in the uterus**

## ACKNOWLEDGEMENTS

We thank Yaoyi Xing, Emeka Okorie, Julia Kelso, Elissa Cosgrove, Nora Brown, Sarah Allen, the Imaging Facility at the Cornell Institute of Biotechnology (RRID:SCR_021741), the Cornell Statistical Consulting Unit and members of the Clark and Wolfner laboratories for advice and assistance with experiments or analysis. We thank Susan Suarez for feedback and advice on the experiments and manuscript.

Stocks obtained from the Bloomington Drosophila Stock Center (National Institutes of Health grant P40OD018537) were used in this study. This work was supported by funds from the National Institutes of Health (R01-HD059060 to A. Clark and M. Wolfner and R01-GM119125 to A. Clark and D. Barbash) and a seed grant from the Cornell Center for Vertebrate Genomics.

## SUPPLEMENTARY MATERIALS

**Table S1.** Drosophila Stocks.

| Full Genotype | Source | Notes |
| --- | --- | --- |
| y[1] w[*] P{y[+t7.7]=nos-phiC31\int.NLS}X; PBac{y[+]-attP-9A}VK00027 | BL #35569 | phiC31/attP9A line used to make guide-expressing flies, from Rainbow Transgenics |
| y[1] w[*]; wg[Sp-1]/CyO; Dr[1]/TM3, Sb[1] | BL #59967 | for balancing transformants |
| w[*]; TM3,Ser /TM6B, Tb | Wolfner Lab Stocks | for balancing transformants |
| w[1118]/Tl{Disc\RFP[tdTom.3xP3]=attP}AByG | BL #78567 | labelled Y chromosome |
| C(1;Y)3, In(1)FM7, v[Of]/C(1)M4, y[1]/O | BL #988 | Source of compound X<sup>X</sup> |
| C(1;Y)1, y[1]/O; ry[506] | BL #4408 | Source of compound X<sup>Y</sup> |
| w[*]; {U6:3-gRNA 3xWDY, w+}attP-9A / TM3 | this study | transformant, expresses *WDY* and *ebony* guides ubiquitously |
| w Protamine-GFP, w+ ;; Bam-Gal4, UAS-Dicer | Chen and McKearin 2003, Manier et al 2010 | Combined into one stock by Caroline Sartain |
| y[1] w[118] ; {nos-Cas9}attP2 / TM3 | Kondo et al 2020 | nos-Cas9 on III |
| yw ; {UAS-WDY RNAi}attP | VDRC_109045 | WDY RNAi Line |
| yw ; {UAS-kl3 RNAi}attP | VDRC_109151 | Kl-3 RNAi line |
| yw ; attP VIE-260B | VDRC_60100 | RNAi Control Vie260 |
| C(1;Y)1, y[1]/C(1)M4, y[1]/Tl{Disc\RFP[tdTom.3xP3]=attP}AByG, WDY[F8] | this study | WDY F8 allele |
| C(1;Y)1, y[1]/C(1)M4, y[1]/Tl{Disc\RFP[tdTom.3xP3]=attP}AByG, | this study | WDY C104 allele |
| C(1;Y)1, y[1]/C(1)M4, y[1]/Tl{Disc\RFP[tdTom.3xP3]=attP}AByG, WDY[C3] | this study | WDY C3 allele |
| C(1;Y)1, y[1]/C(1)M4, y[1]/Tl{Disc\RFP[tdTom.3xP3]=attP}AByG, [C7] | this study | Control C7 allele |
| C(1;Y)1, y[1]/C(1)M4, y[1]/Tl{Disc\RFP[tdTom.3xP3]=attP}AByG, [G107] | this study | Control G107 allele |

**Table S2.**
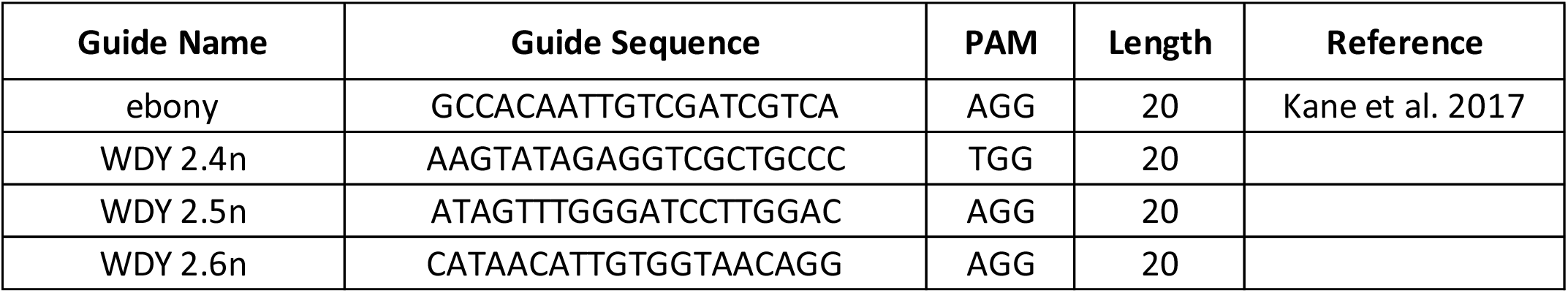
Guide Sequences.

**Table S3.**
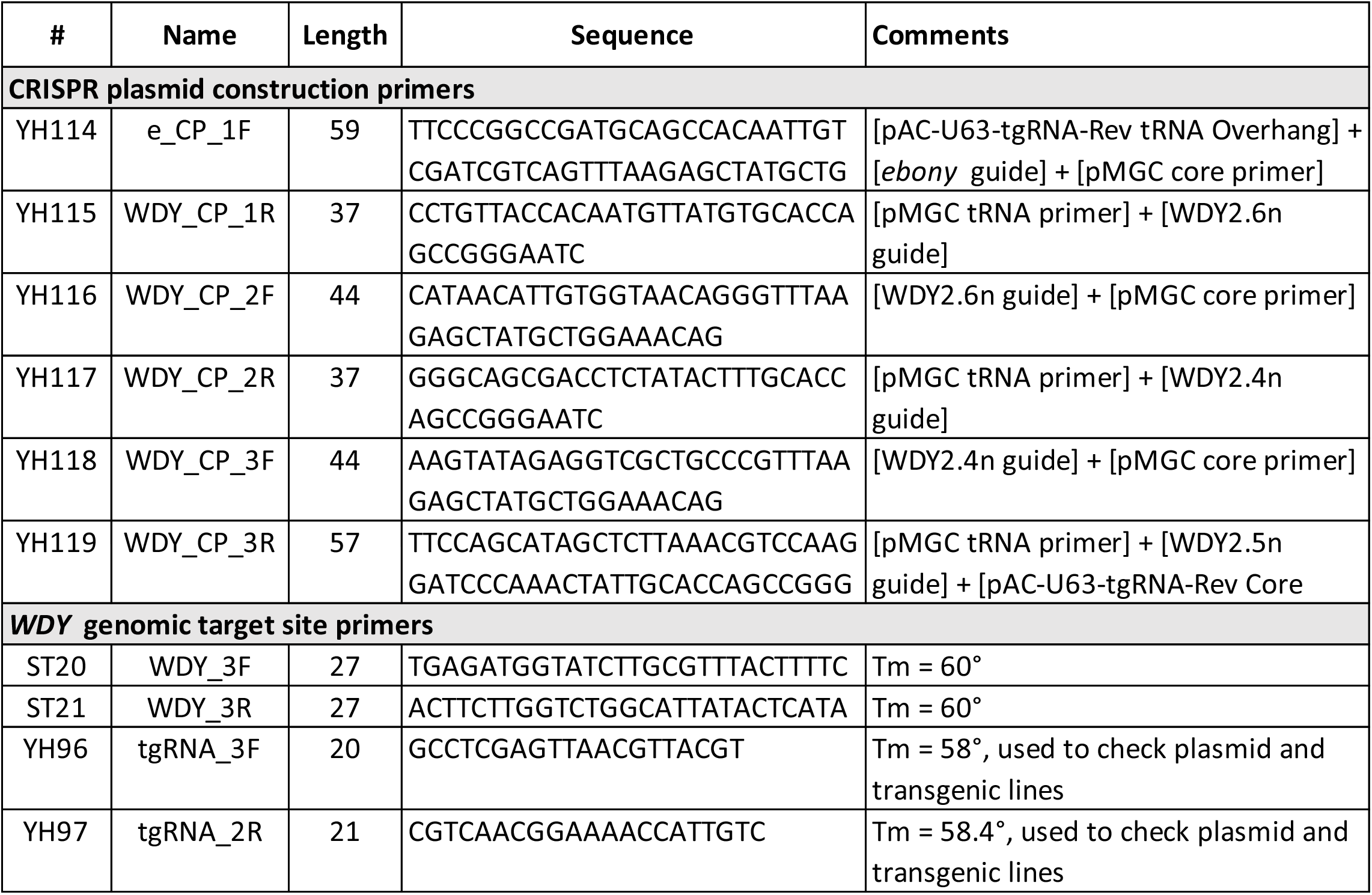
Primer Sequences.

**Table S4.** Alleles Generated.

| Allele | Characteristics of the allele |
| --- | --- |
| Y_Tomato | unedited marked Y chromosome |
| Control_G107 | went through crossing scheme but showed no changes at target site |
| Control_C7 | went through crossing scheme but showed no changes at target site |
| WDY[F8] | 547bp deletion between guides 2.4n-2.5n |
| WDY[C104] | 545bp deletion between guides 2.4n-2.5n |
| WDY[C3] | 443bp deletion between guides 2.6n-2.5n + 3bp change at guide 2.4n |

**Table S5.** Sterility Test of *WDY*.

| Genotype | Allele | Fertile | Sterile |
| --- | --- | --- | --- |
| Control | tdTomato | 18 | 0 |
| Control | C7 | 17 | 0 |
| Control | G107 | 17 | 0 |
| WDY | F8 | 0 | 19 |
| WDY | C104 | 0 | 19 |
| WDY | C3 | 0 | 19 |
| Control | RNAi | 18 | 1 |
| WDY | RNAi | 5 | 15 |
| kl-3 | RNAi | 0 | 10 |

**Table S6.**
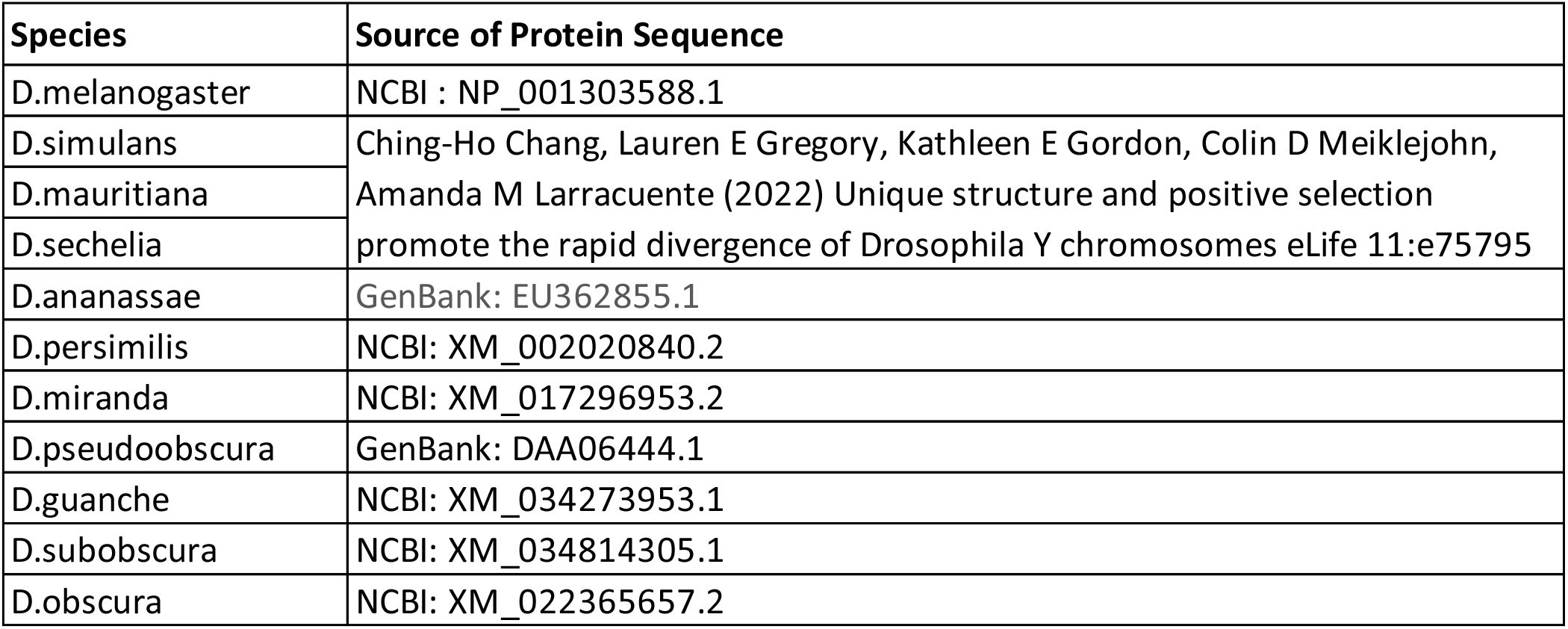
WDY Orthologue Sequences.

| Species | Source of Protein Sequence |
| --- | --- |
| *D.melanogaster* | NCBI : NP_001303588.1 |
| *D.simulans* | Ching-Ho Chang, Lauren E Gregory, Kathleen E Gordon, Colin D Meiklejohn, Amanda M Larracuente (2022) Unique structure and positive selection promote the rapid divergence of *Drosophila* Y chromosomes eLife 11:e75795 |
| *D.mauritiana* |  |
| *D.sechelia* |  |
| *D.ananassae* | GenBank: EU362855.1 |
| *D.persimilis* | NCBI: XM_002020840.2 |
| *D.miranda* | NCBI: XM_017296953.2 |
| *D.pseudoobscura* | GenBank: DAA06444.1 |
| *D.guanche* | NCBI: XM_034273953.1 |
| *D.subobscura* | NCBI: XM_034814305.1 |
| *D.obscura* | NCBI: XM_022365657.2 |

**Figure S1.**
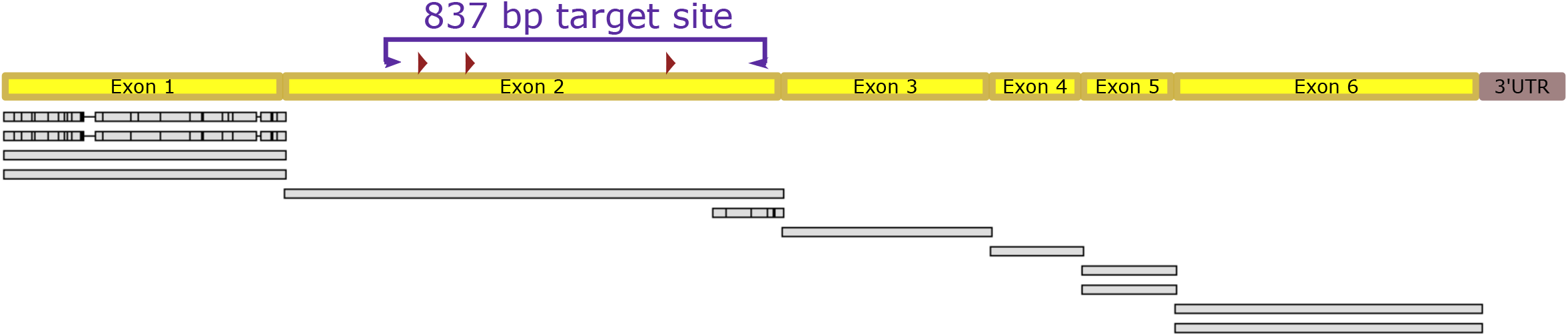
CRISPR target site on exon 2 of *WDY* had no identifiable duplicates. Exon structure of *WDY* (introns not shown) with CRISPR target site marked. Purple arrows indicate primers, red triangles indicate guides. Additional copies of *WDY* regions on the Y chromosome are displayed as grey bars underneath the *WDY* schematic.

**Figure S2.**
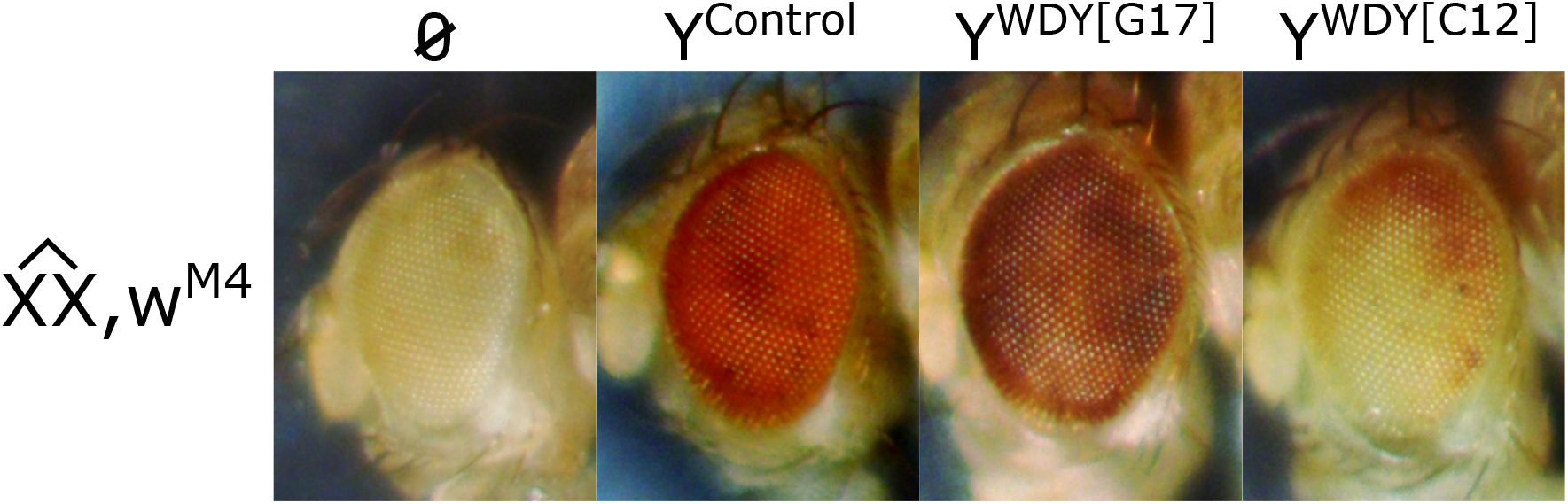
Position effect variegation observed in some lines after CRISPR editing at *WDY*. Adult eyes of females with compound-X containing the position effect variegation marker, w^M4^, and no Y chromosome (A), an unedited Y chromosome (B), or CRISPR-edited Y chromosomes G17 (C) or C12 (D).

**Figure S3.**
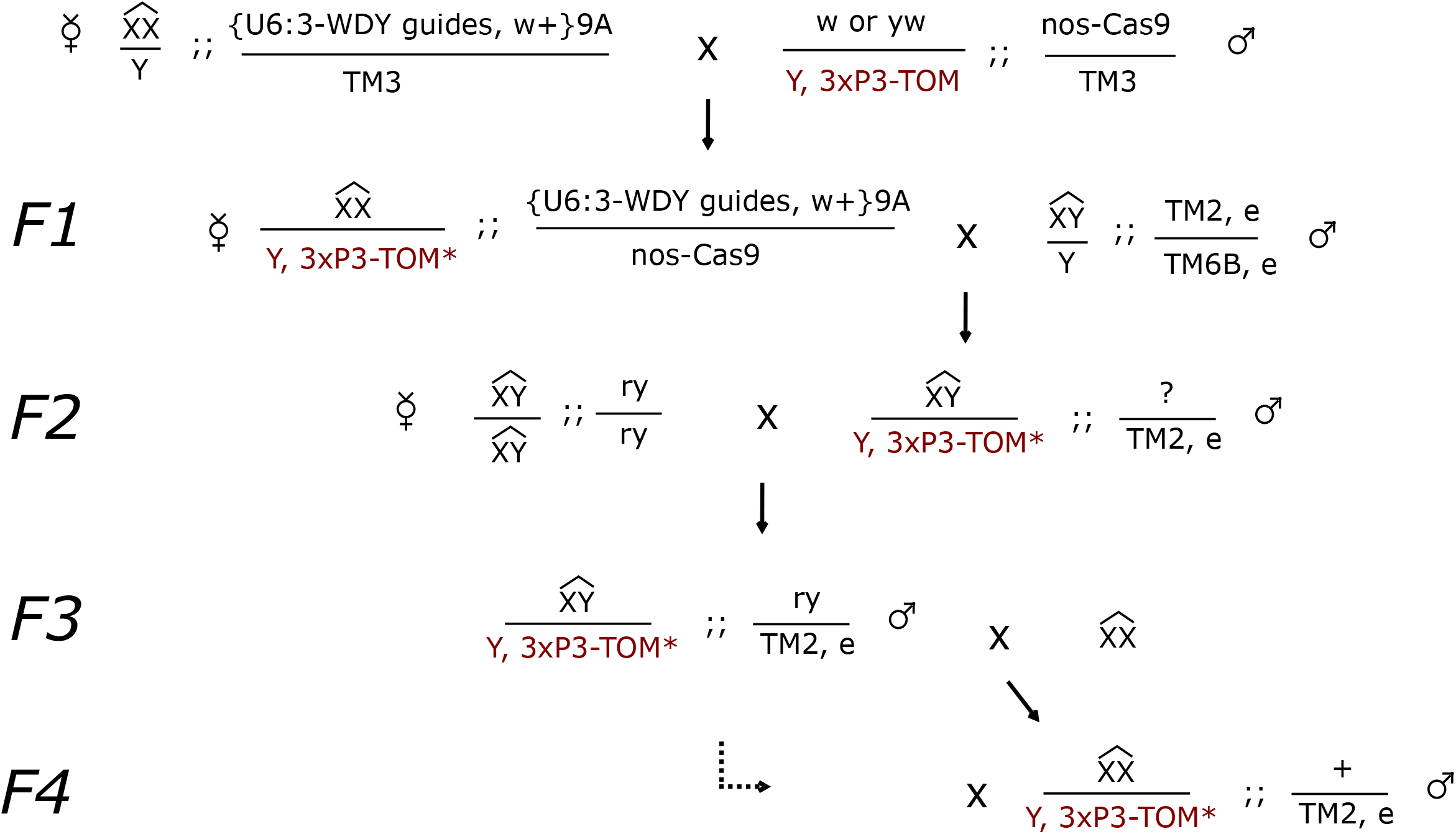
Crossing scheme for inducing CRISPR, then balancing the edited Y chromosome using compound-X and attached X-Y chromosomes.

**Figure S4.**
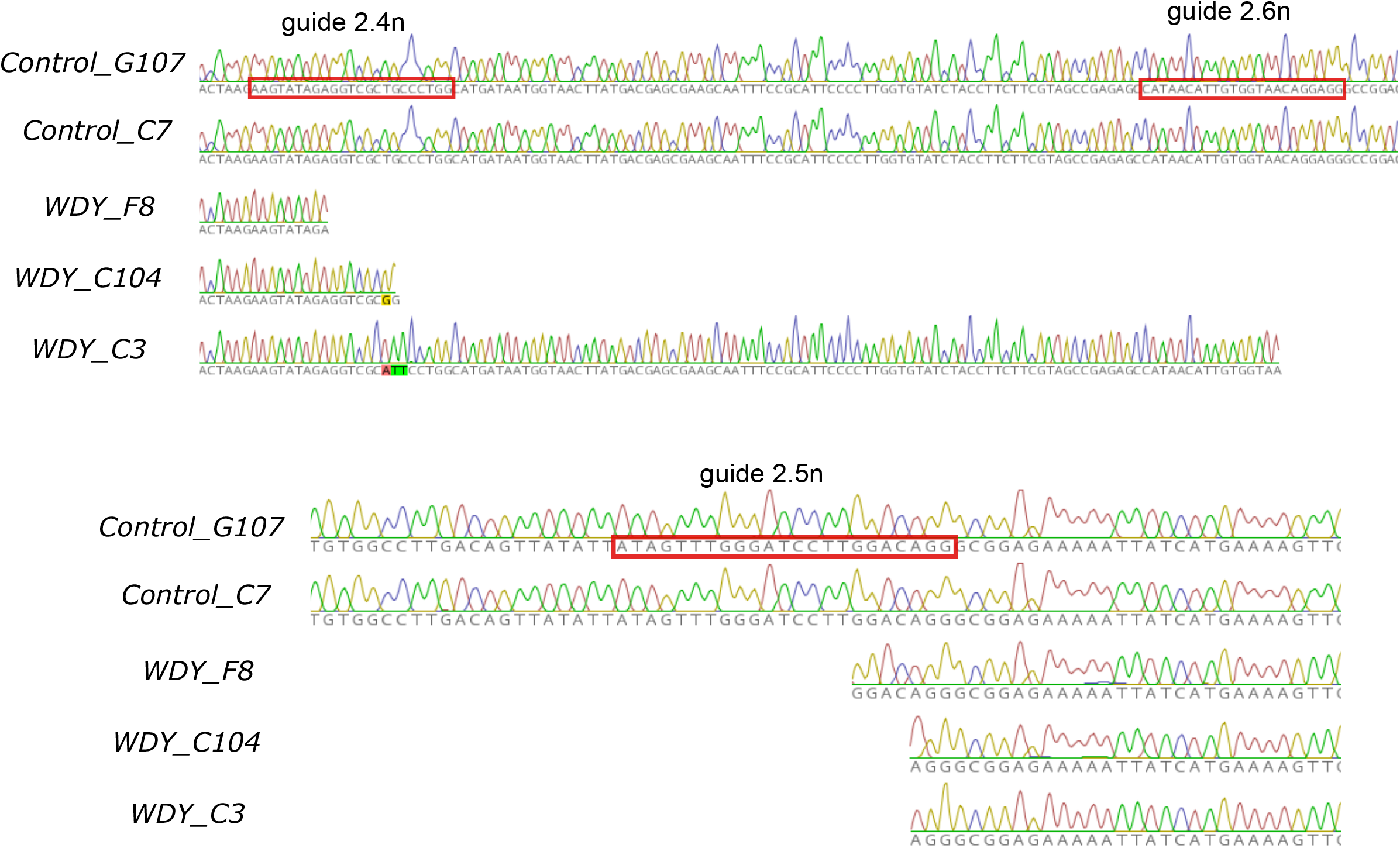
Sequencing results in target region for *WDY* and control alleles used in this study. Guide sites are marked with red boxes. Nucleotide changes are highlighted. Gaps represent deletions.

**Figure S5.**
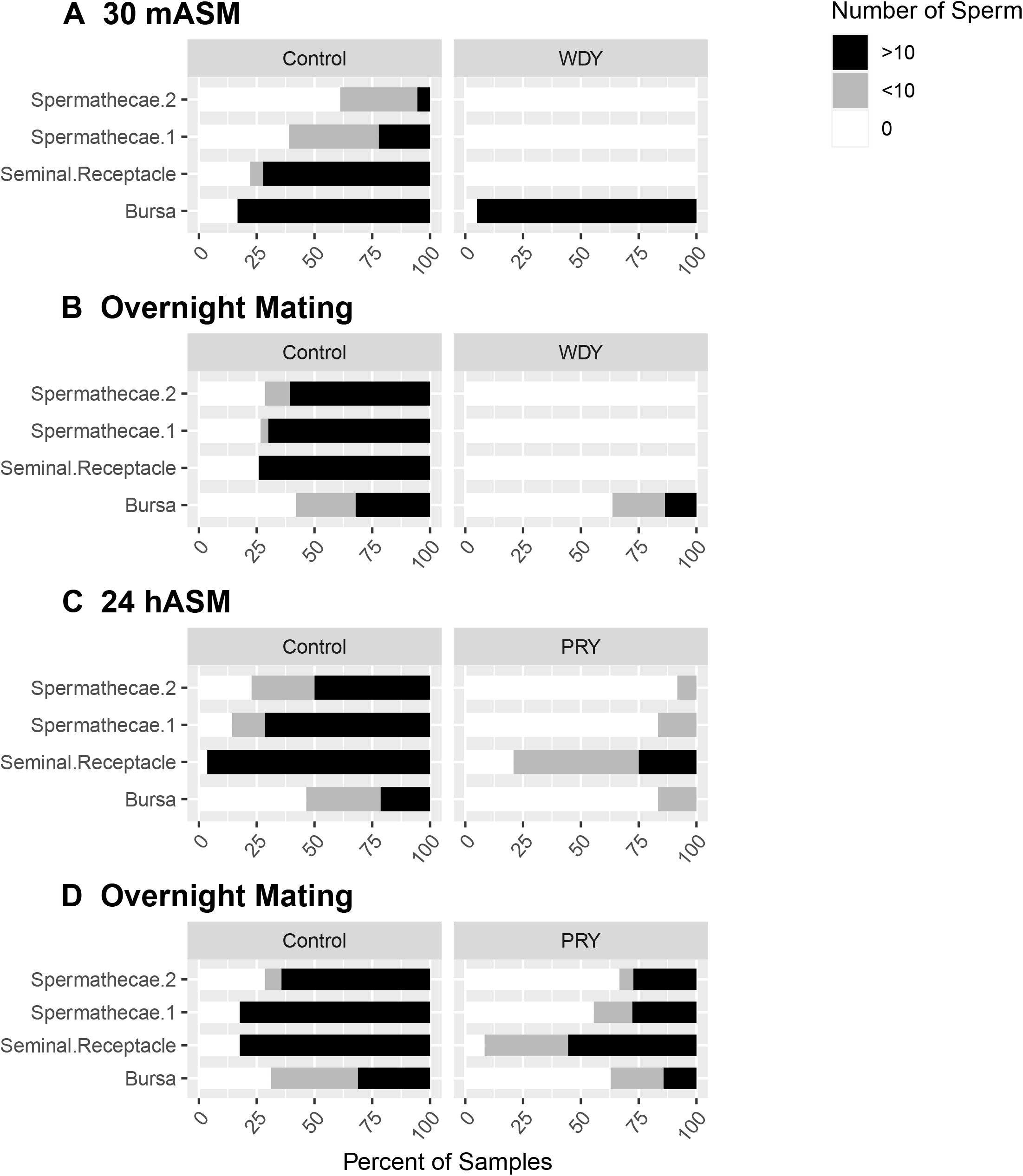
Distribution of *WDY* and *PRY* mutant sperm in the female reproductive tract after mating. (A) Quantification of *WDY* mutant sperm’s distribution in the female reproductive tract 30 mASM. (B) Quantification of *WDY* mutant sperm’s distribution in the female reproductive tract after overnight mating. (C) Quantification of *PRY* mutant sperm’s distribution in the female reproductive tract 24 hASM. (D) Quantification of *PRY* mutant sperm’s distribution in the female reproductive tract after overnight mating.

## METHODS

### Drosophila Stocks and Husbandry

Flies were reared on a cornmeal-agar-sucrose medium (recipe available at https://cornellfly.wordpress.com/s-food/) at 25°, with a 12 hr light-dark cycle. The stocks used in this study are described in Table S1.

### Generation of a *WDY* Mutant with CRISPR

Three 20 base pair guide RNAs were designed to target exon 2 of *WDY*, a region of the gene with no known duplications (Figure S1) ^1^. We also targeted *ebony*, a visible Co-CRISPR marker ^2^. Guide sequences were incorporated into pAC-U63-tgRNA-Rev (Addgene, Plasmid #112811) which is analogous to the “tgFE” construct from [3]. This was done by appending guide RNA sequences to tracrRNA core and tRNA sequences from pMGC (Addgene, Plasmid #112812) through tailed primers (Table S3) to create inserts that were then inserted by Gibson Assembly into a *SapI*-digested pAC-U63-tgRNA-Rev (Addgene, Plasmid #112811). The plasmid backbone contained attB, and we used Phi-C31 to integrate it into an attP-9A site on chromosome 3R. The construct was injected into *yw nanos-phiC31; PBac{y+-attP-9A}VK00027* by Rainbow Transgenic Flies Inc. Transformants were identified by eye color from a *mini-white*+ marker. Transformants express the four guides ubiquitously under the U6:3 promoter as a single polycistronic transcript that is processed by the endogenous cellular tRNA processing machinery (RNase P and Z) to release the individual mature gRNAs and interspersed tRNAs. The transformants were balanced and inserts were confirmed by PCR and sequencing. A few of the transformants had light-red eyes, but we only used those with dark-red eyes.

Different combinations of transformants and germline Cas-9 drivers were tested for editing efficiency (data not shown). Males containing *vasa-Cas9* and our guide RNAs were sterile, while females showed a 6.2% CRISPR efficiency, based on the generation of *ebony* mutants. In contrast, F1 males from crosses with *nanos-Cas9* drivers on chromosomes 2 and 3 produced progeny. F2 progeny (from male and female crosses combined) showed that these two lines had editing efficiencies of 2.2% and 2.6%, respectively. Our observation of higher efficiency and sterility from *vasa-Cas9* is consistent with the earlier ^4^ and higher somatic ^5,6^ protein expression of Vasa versus Nanos, as well as the RNAi phenotype of *WDY* ^7^. We proceeded to make stable mutants by crossing transformant #1 to *nanos-Cas9* in a compound chromosome background.

Our crossing scheme for creating *WDY* alleles is shown in Figure S3. We combined the *nanos-Cas9* driver on chromosome 3 with a Y chromosome marked with *3xP3-tdTomato* ^8^. We also combined the guide-expressing insert on 3R with a compound 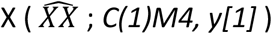. CRISPR editing occurred in F1 females 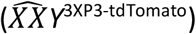 that carried the marked Y chromosome. By crossing to a compound 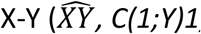 we were able to establish balanced lines from 55 *ebony* and 5 non-ebony F2 flies. Males were of genotype 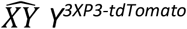 and were fertile regardless of CRISPR-mediated edits of the free Y. We screened these lines for visible deletions in the *WDY* target site – first by gel, then by sequencing. Alleles derived from our crossing scheme are listed in Table S4 and described in Figure S4. They are maintained as stable lines with 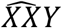 females and 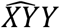 males; the free Y chromosome is edited.

In several of our lines, we saw varying, intermediate degrees of position effect variegation (PEV) (Figure S2). This corresponded with either failed amplification at the target site in *WDY* or the presence of several bands of unexpected size. Based on our previous results when editing *FDY* with CRISPR^7^, we hypothesized that these mutations were large deletions in the Y chromosome, and thus did not phenotype these mutants for sperm or fertility characteristics. C(1)M4 contains *white*[*mottled-4*], a PEV marker that is highly sensitive to Y-chromosome dosage. 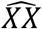 females with C(1)M4 have mostly white eyes, while 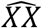 females have an almost entirely red eye. We previously showed that lines with visibly altered PEV lacked large sections of the Y chromosome^7^. Such deletions may be caused by the presence of uncharacterized copies of the target region present in unassembled regions of the Y chromosome.

### Sterility, Mating, and Sperm Storage

Crosses and experiments were done with flies 2-5 days after eclosion (dAE). To test for sterility, we crossed individual XY males to 4 Canton S virgin females in a food containing vial with wet yeast. Adults were transferred to a new vial after one week. Crosses were scored for the presence of progeny. 15-20 crosses were tested per line. For experiments that required timing from the start of mating, one Canton S virgin female was mated to three males of a given genotype and flies were observed. Once mating began the time was noted. Females were analyzed or flash frozen in liquid nitrogen for 30 minutes, 2 hours, or 24 hours after the start of mating (30 mASM, 2 hASM, 24 hASM). Reproductive tracts were dissected from frozen females in PBS, fixed in 4% paraformaldehyde, and mounted in Vectashield with DAPI. Samples were imaged on an Echo Revolve microscope or a Leica DMRE confocal microscope.

### Sperm Counting with Imaris software

To quantify sperm transferred, female reproductive tracts 30mASM were imaged on a Leica DMRE confocal using standardized settings. Two μm Z-stacks through each sample were collected. Using Imaris 9.8.0 software (RRID:SCR_007370), first the female reproductive tract was extracted in each image by manually drawing a contour surface. The mating plug and cuticle were specifically excluded due to their high autofluorescence ^9^ Second, protamine-labelled sperm heads were automatically detected using the “Surfaces” function (smoothing and background elimination enabled, 2.0 µm surface grain size, 1 µm diameter of largest sphere, 2.747-13.048 manual threshold, >15 quality, <0.8 sphericity). Counts of transferred sperm from control and *WDY* males were statistically compared using a Student’s t-test in R software.

### Sperm-tail Beat Frequency Analysis

Tail-beat frequency was measured for sperm dissected from the reproductive tracts of males 2-5dAE or females 2-5 dAE and 30 mASM into PBS. Sperm were released into a 15 ul drop of PBS on a glass slide by tearing the male seminal vesicle or female uterus. Sperm were observed under brightfield optics with an Olympus BX51WDI microscope and a 50x LMPLFLN objective. Eight second raw movie clips at 1280×720 resolution and 60 frames per second were captured from 4-6 different regions around the sperm mass using a Canon EOS Rebel T6 camera. Dissected sperm masses all contain sperm tails beating at a range of frequencies - we specifically quantified the beat frequency of the 1-2 fastest-beating sperm tails from each clip.

To measure the sperm tail-beating frequency, video clips were imported to FIJI (RRID:SCR_002285) using the ffmpeg plug-in. From each clip, we measured beat frequencies of the 1-2 fastest-beating sperm, limited to tails that were not overlapping or entangled with other sperm tails. A selection line was drawn across an isolated section of sperm tail. A 1-pixel “Multi Kymograph” was generated which shows pixel intensities across the selection line on the X-axis for each frame along the Y-axis. The beating of the sperm tail appears as a traveling wave form. The number of beats and the number of frames were counted for the region where the sperm tail remained in focus and isolated from other tails. Beat frequency was then calculated as: Hz = (# beats X 60 fps) / # frames. Ten measurements were made per individual fly. Approximately one-third of samples were scored blind, and statistical analysis indicated consistent results whether samples were scored blind or not.

Sperm tail-beat frequencies were measured from a minimum of three individuals of each allele. Using the lme4 package in R, linear mixed models were fitted to the data, incorporating the individual as a random effect and experimental batch and the experimenter who measured beat frequency as fixed effects. We then ran a Likelihood-Ratio test to compare the model with and without “Genotype” as a fixed effect.

### Sperm Swimming Analysis

Videos of sperm swimming were acquired from either male reproductive tracts or female reproductive tracts 1 hASM. Tracts were dissected and mounted in 15 µL PBS. Spacers (2 layers of double-stick tape) were used to avoid compression of the tissue by the coverslip. Fluorescent sperm heads were recorded through screen recording of the preview window on an Echo Revolve. We used ffmpeg (RRID:SCR_016075) to convert videos to constant frame rate of 60 fps and .mov format. Videos were then imported into FIJI (RRID:SCR_002285) using the ffmpeg plugin. We manually tracked sperm heads across 60 frames using the “Manual Tracking” plugin in FIJI. The tracking shown in Figure 3 represent movement across 30 frames.

### WDY Annotation and Sequence Comparisons

EF hand motifs were identified by searching (using Geneious software, RRID:SCR_010519) for the canonical and pseudo PROSITE motif consensus sequences defined in [^10^] and allowing for a maximum 1 base pair mismatch. Because the pseudo-EF hand motif contains a variable size region, there were two potential start locations in the sequence – residue 44 or 47. However, the Alphafold prediction showed residues that should form the loop region would instead form part of the alpha helix in the motif beginning at residue 44. We therefore favored the motif beginning at residue 47. Locations of the calcium binding residues were determined based on the consensus sequence logograms in [^10^].

Three WD40 domains ^11^ were originally identified in the protein sequence based on homology. Flybase reported a handful of WD40 repeats (2 for Pfam and 8 for SMART) were identified. 4 to 16 of these repeat domains may together form a circular beta propeller structure called a WD40 domain ^12–14^; however, insufficient WD40 repeats were identified in WDY to predict the presence of a WD40 domain. We used the a structural prediction of *D. melanogaster* WDY by Alphafold (PDB B4F7L9) ^15^ to identify the locations of the characteristic β-propeller, consisting of 4 antiparallel sheets ^13^. WDY is predicted to form two WD40 domains - one with 6 WD40 repeats and one with 7 WD40 repeats.

WDY ortholog sequences were obtained as described in Table S6. The proteins were aligned in Geneious using a BLOSUM cost matrix with a gap open cost of 10 and a gap extend cost of 0.1.

